# RecBCD Prevents Bacterial Autoimmunity by Clearing Self-DNA Ends Produced During Replication Termination

**DOI:** 10.64898/2026.09.30.755081

**Authors:** Apurva A. Govande, Brooke Ramsey, Irem Unlu, Susan K. Amundsen, Toni A. Nagy, Albur Hassan, Marina Smiley, Catherine Trautman, Aaron T. Whiteley, Gerald R. Smith, Sangjin Kim, Hoong Chuin Lim

## Abstract

Viral-induced DNA damage triggers both eukaryotic and prokaryotic innate immunity. Here, we report that self-DNA ends generated routinely during normal termination of bacterial replication suffice to trigger the P2-encoded OLD antiphage system. Normally, these physiological triggers are cleared by the dual helicase-nuclease activities of the conserved RecBCD enzyme, which acts as an immune safeguard analogous to mammalian TREX1 exonuclease and plant DBR1 debranching enzyme. However, when this clearance fails, the transient self-DNA ends persist and are recognized by ATP-bound OLD dimers via a positively charged channel across their ATPase subunits. ATP hydrolysis then activates OLD to cleave tRNAs, causing cytotoxicity. Single-cell microscopy reveals that dozens of OLD molecules associate with kilobases of adjacent DNA, amplifying the sparse physiological DNA end signal. Our findings reveal how an endogenous byproduct poses a constant autoimmune threat in bacteria, highlighting a universal molecular conflict across kingdoms and the convergent solutions evolved to resolve it.

## Introduction

Immune systems targeting invading nucleic acids must reliably distinguish invading nucleic acids from chemically identical self-DNA and RNA^1–3^. Systems like CRISPR-Cas and Argonaute enforce strict base-pairing rules to discriminate against self-nucleic acids^1,4,5^. Restriction-modification systems and eukaryotic RIG-I recognize modifications and structural motifs^1,6^, and systems like bacterial SNIPE and mammalian cGAS rely on spatial regulation^7–9^. However, to ensure avoidance of activation by self-nucleic acids, eukaryotes recruit housekeeping nucleases as a sanitization layer to clear immunogenic self-nucleic acids^2,9^. In mammals, mutations in nucleases such as TREX1 cause nuclear DNA to accumulate in the cytoplasm, triggering cGAS-STING autoimmunity^10,11^. In plants, loss of the RNA debranching enzyme DBR1 leads to intron lariat accumulation and aberrant activation of small RNA-based defenses12. Although chemical- and phage-induced DNA damages are known to trigger bacterial defense systems^13–15^, whether bacteria deploy analogous mechanisms to suppress immunogenic endogenous nucleic acids during normal growth is unknown.

A hint that such mechanisms exist lies in the unexplained behavior of “RecBCD guards”— antiphage systems like OLD, Retron Ec48, PARIS, and Gabija that monitor the conserved host RecBCD helicase-nuclease^16–20^. Because phages frequently manipulate RecBCD to favor their own recombination and replication pathways, these guards trigger defense upon sensing RecBCD perturbation. Interestingly, many RecBCD guards become toxic in uninfected *recBCD* mutants^16,17,20,21^, suggesting they detect an endogenous immunostimulatory signal that accumulates in the absence of RecBCD. Although this autoimmune phenomenon was first reported in 1969^16^, the identity of this signal and why it arises only when RecBCD is absent have remained elusive.

To address these questions, we focused on the OLD (Overcoming Lysogenization Defect) system, which is deployed by *E. coli* prophage P2 to abort phage λ infection^22^. This single-protein system is the founding member of a widespread family of defense systems combining an ABC-type ATPase and Toprim domain^23–25^ and the first system identified to trigger host toxicity upon RecBCD impairment*^16^*. Despite its long-standing annotation as a DNA nuclease^25,26^, we recently established that OLD drives toxicity by severing the anticodon stems of host tRNAs^27^, a conclusion supported by a recent study^28^. Building on this finding, we show that the toxic tRNA cleavage activity of OLD is triggered by double-stranded DNA (dsDNA) ends that accumulate in cells lacking RecBCD. These DNA intermediates arise routinely during normal replication termination but are rapidly cleared by the RecBCD complex. We define the molecular basis for how OLD senses these ends, reconstitute its activation in vitro, and propose a signal amplification mechanism that enables the sensitive detection of this sparse physiological trigger.

## Results

### RecBCD nuclease activity suppresses OLD-mediated toxicity

Expression of OLD *in trans* severely compromises the viability of *E. coli* mutants lacking any component of the RecBCD complex (Fig. 1a). In contrast, a *recA* mutant is only weakly sensitive, confirming that that OLD toxicity is not caused by a recombination defect^22^. We sought to determine how RecBCD normally suppresses OLD toxicity. Because RecBCD integrates DNA unwinding, recognition of Chi recombination hotspots, and exonuclease activities within a single multifunctional complex^29^, we first sought to determine which of these functions is required to suppress OLD. To do this, we assayed tolerance to OLD expression across a panel of *E. coli* strains carrying chromosomal alleles that selectively impair RecBCD function (Fig. 1a). Chi-site recognition is dispensable for suppressing OLD toxicity, as a strain defective for this function (*recBC*^1004^*D*) tolerates OLD comparably to the wild-type *E. coli*. In contrast, strains encoding nuclease-dead variants such as RecB^K29Q^CD and RecB^K29Q^CD^K177Q^, including those with helicase activity (RecBC [Δ*recD*] and RecB^D1080A^CD), fail to tolerate OLD. OLD expression slightly impaired the growth of a strain encoding RecBCD^K177Q^, which retains ∼20% nuclease activity, indicating partial tolerance. Overexpression of RecBCD^K177Q^ fully restored tolerance in a Δ*recBCD* strain, demonstrating that full suppression of OLD toxicity requires a critical threshold of RecBCD nuclease activity (Fig. 1b). Unexpectedly, overexpression of the nuclease-dead, helicase-active RecBC or RecB^D1080A^CD also partially rescued the OLD-mediated lethality (Fig. 1b). However, unlike the nuclease-active RecBCD or RecBCD^K177Q^, this helicase-mediated suppression strictly depends on the non-cognate single-stranded DNA exonuclease RecJ (Fig. 1c). Collectively, these results indicate that suppression of OLD activation requires both DNA unwinding and DNA cleaving activities. This dual requirement suggests that RecBCD actively eliminates an end-exposed double-stranded DNA (dsDNA) intermediate that would otherwise activate OLD. Supporting this interpretation, OLD toxicity can be partially suppressed by an *sbcA* mutation, which activates the RecET system to act on linear DNA ends (Fig. S1).

**Figure 1.**
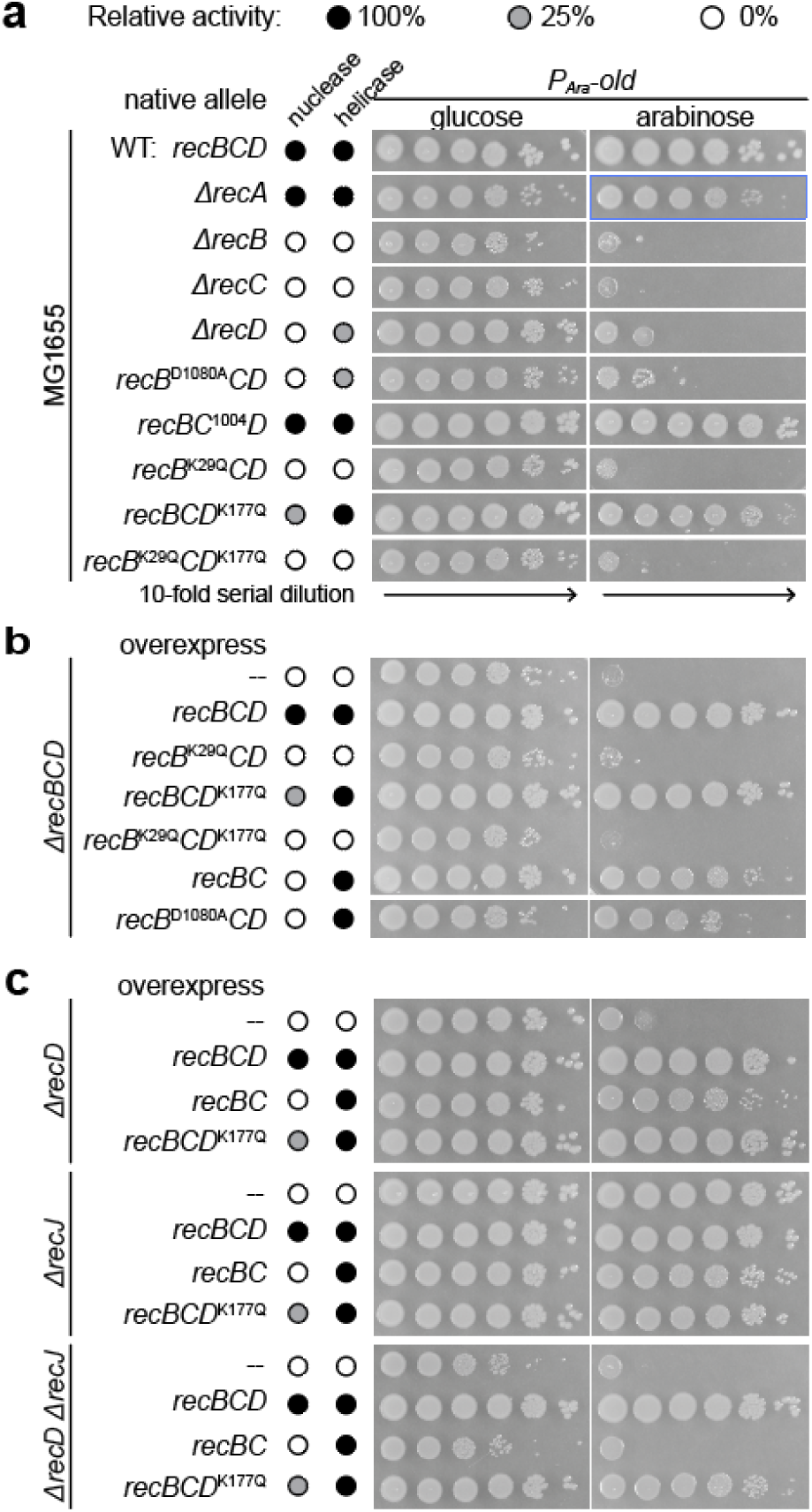
Both the helicase and nuclease functions of RecBCD are required to suppress OLD toxicity. **(a)** Ten-fold serial dilutions of cultures of *E. coli* MG1655 strain or its derivatives carrying the indicated *recBCD* alleles at the native chromosomal locus. These strains harbor an expression plasmid producing OLD from the *P_BAD_* arabinose-inducible promoter. The dilutions were spotted on LB medium agar supplemented with either glucose or arabinose to repress or induce the production of OLD as indicated. **(b, c)** Same as **a**, but with the indicated *recBCD* alleles constitutively produced from a second plasmid in the specified mutant backgrounds.

### OLD localizes to uncleared DNA ends at bacterial host chromosome termini

We next sought to determine how this putative DNA trigger is generated, and whether it possesses specific sequence or structural features. We reasoned that the triggering DNA likely binds OLD directly, meaning its association with the protein should temporally track the kinetics of activation. To capture this dynamic, we initiated RecBCD inactivation by expressing the λ-encoded inhibitor Gam^30^ in wild-type cells and performed ChIP-seq using a functional, C-terminally FLAG-tagged OLD variant (OLD-F) (Fig. S2a and Fig. 2a, b). Despite some background, OLD-F immune-precipitation reproducibly enriched a ~20 kb region encompassing the replication terminus within 20 minutes of Gam induction, and this signal intensified by 40 minutes (Fig. 2c and Fig. S2b), identifying this locus as a candidate source of the DNA trigger. Terminus recruitment required ATP-induced conformational changes of OLD, but not its tRNA cleavage or ATP hydrolysis activities (Fig. 2d). The tRNA-cleavage deficient mutant (OLD^E402A^-F) and the ATP-hydrolysis-deficient Walker B mutant (OLD^E300Q^-F) localized to the terminus normally^27,31^. In contrast, the OLD H-loop mutation (H332A)^31^ —which similarly permits ATP binding but not hydrolysis—completely abolished recruitment. In a related ABC-type ATPase protein HylB, the equivalent histidine stabilizes the transition state required for ATP hydrolysis and mediates intersubunit communication within the ATPase dimer^32^, suggesting that ATP binding triggers a structural rearrangement to enable target engagement. Interestingly, ChIP-seq signals of OLD variants with intact ATPase peaked immediately adjacent to *dif* and *TerC* (Fig. 2d). In contrast, the corresponding ‘peaks’ of ATP hydrolysis-defective variant (OLD^E300Q^) migrated away from those sites (Fig. 2d and Fig. S2d), indicating that ATP turnover regulates the dynamics of DNA engagement.

**Figure 2.**
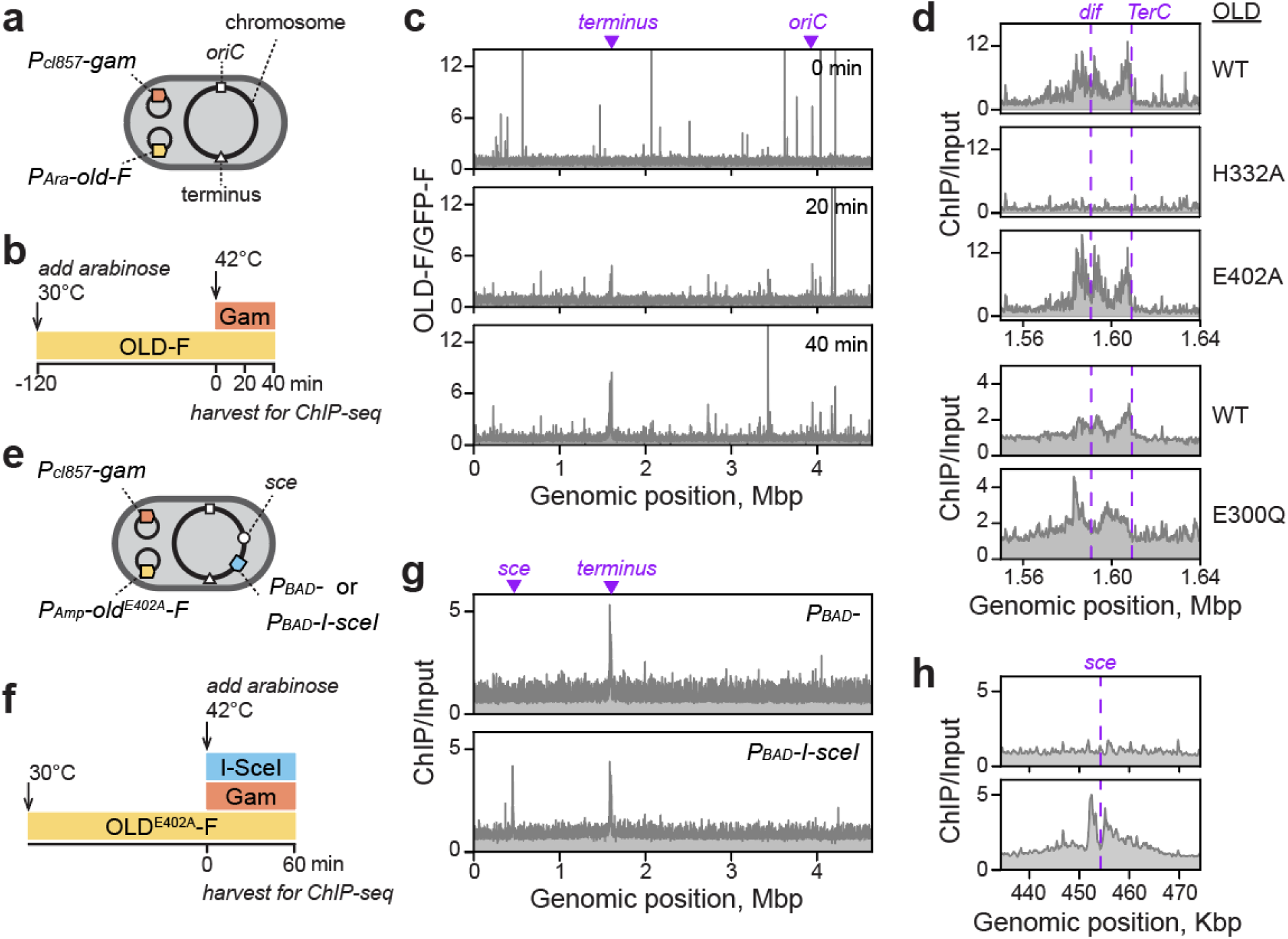
ATP-bound OLD binds to self-DNA ends at the bacterial replication terminus upon RecBCD inhibition. **(a, b)** Scheme for OLD-F and Gam induction prior to ChIP-seq. MG1655 carrying the indicated plasmids were cultured at 30°C in the presence of arabinose to induce OLD-F expression for 2 hours before shifting to 42°C to induce Gam production. Cells were crosslinked at 0-, 20-, and 40-minute (min) time points for ChIP-seq. **(c)** Representative genome-wide ChIP-seq profiles of OLD-F collected at the indicated time points post-Gam induction. Signals were normalized to GFP-F to reduce background signal common to both OLD-F and GFP-F samples. Input-normalized profiles for OLD-F and GFP-F are shown in Fig. S2b. **(d)** Magnified view of ChIP-seq profiles near *dif* and *TerC* at 40 min post-Gam induction. Genome-wide profiles are shown in Fig. S2c. **(e, f)** Experimental setup for targeted DSB induction prior to ChIP-seq. The experimental strain harbors an ectopic *sce* site at position 454244 in the MG1655 genome and either *P_BAD_*-*I*-*sceI* or an empty *P_BAD_* control at the λ attachment site. OLD^E402A^-F and Gam were produced in trans from the constitutive *P_AMP_* and *P_cI857_*, respectively. The cells were crosslinked 60 minutes after arabinose addition (to induce I-*Sce*I) and a temperature shift to 42 °C (to induce Gam) for ChIP-seq. **(g)** Representative genome-wide ChIP-seq profiles of OLD^E402A^−F 60 minutes following concurrent induction of Gam and I-*Sce*I. **(h)** Magnified view of ChIP-seq profiles shown in **(g)** highlighting the genomic region surrounding *sce*. Data are representative of two biological replicates.

The localization pattern of OLD suggested a connection to the aberrant replication termination characteristic of *recBCD* mutants, which frequently leaves sister chromosomes physically tethered at this site^33,34^. In *recB* and *recC* mutants, subsequent cell division across these unresolved chromosomes mechanically severs the DNA, triggering the loss of hundreds of kilobases of terminus DNA^33–36^. However, in the *recD* mutant—which is also sensitive to OLD (Fig. 1a)—this region is over-replicated rather than degraded^34^. We therefore hypothesized that OLD senses an early DNA intermediate common to all three *recBCD* mutants before further processing leads to different outcomes. Supporting this, our OLD ChIP signal localized to a comparatively narrow region featuring three prominent peaks—two flanking *dif* and a third abutting *Ter*C (Fig. 2d). Notably, both sites are hotspots for distinct types of dsDNA breaks (DSBs) in wild-type *E. coli*^33^. DSBs at *dif* are two-ended, arising when XerCD-mediated cleavage occasionally fails to re-seal^33^. In contrast, DSBs at *Ter* sites are one-ended, resulting from the directional arrest of incoming replication forks^33,37^. When a second co-directional replication fork arrives at this barrier before an opposing fork can fuse with the first, a one-ended DSB forms on the blocking face^33,37^—precisely where we observed the *Ter*C-adjacent ChIP-seq signal (Fig. 2d). Because DNA ends at these sites are normally processed by RecBCD and accumulate in its absence^33^, they represent compelling candidates for the signal that recruits OLD.

To test this model, we inserted an ectopic I-*Sce*I cleavage site (*sce*) midway between the origin (*oriC*) and terminus in a strain chromosomally encoding an inducible I-*Sce*I nuclease^33^ (Fig. 2e). Because OLD-mediated tRNA cleavage might suppress the expression of I-*Sce*I nuclease itself, we constitutively expressed a nuclease-dead OLD^E402A^-F that still localizes to the terminus normally (Fig. 2d). As expected, OLD^E402A^-F accumulated at the chromosomal terminus upon Gam-mediated RecBCD inhibition in a control strain that contains the *sce* site but does not encode I-*Sce*I (Fig. 2f, g). However, co-expression of I-*Sce*I and Gam resulted in an additional signal consisting of two peaks flanking *sce* (Fig. 2g, h), indicating that OLD localizes to both ends of the DSB. Collectively, these results indicate that OLD acts as a dsDNA-end sensor, detecting persistent RecBCD substrates as a readout for enzyme deficiency.

### DNA-end recognition by an ATPase channel allosterically activates tRNA cleavage

Next, to determine whether DNA-end recognition activates OLD, we sought to disrupt this interaction and ask whether it abolishes toxicity in RecBCD-deficient cells. Structural modeling predicts that OLD forms a butterfly-shaped OLD dimer in complex with two ATP molecules and a 30-nt DNA duplex (Fig. 3a). The ATPase domains drive this dimerization, creating two composite ATP-binding sites at the subunit interface (Fig. S3a). Dimerization also creates a positively charged DNA-binding channel across the top of the ATPase domains (Fig. 3a, b), distant from the Toprim domains (Fig. 3a). Consistent with an evolutionary link between the two ATPases^24^, this predicted binding mode mirrors how the eukaryotic DNA damage-sensor Rad50 engages with DNA ends^38^ (Fig. S3b). Guided by this model, we mutated three clustered lysines (K159, K161 and K168) to alanines to selectively disrupt DNA-end binding (Fig. 3c). Supporting this channel acting as the DNA sensor, ChIP-seq analysis showed that this variant (OLD^Patch1^) failed to enrich at the replication terminus upon RecBCD inhibition (Fig. 3d). Moreover, OLD^Patch1^ and two additional channel mutants (OLD^Patch2^ and OLD^Patch3^) became completely non-toxic during RecBCD inhibition (Fig. 3e and Fig. S3c) and failed to block λ phage replication (Fig. S3d). Combined, these results functionally link DNA-end recognition to the activation of OLD’s tRNA cleavage activity.

**Figure 3.**
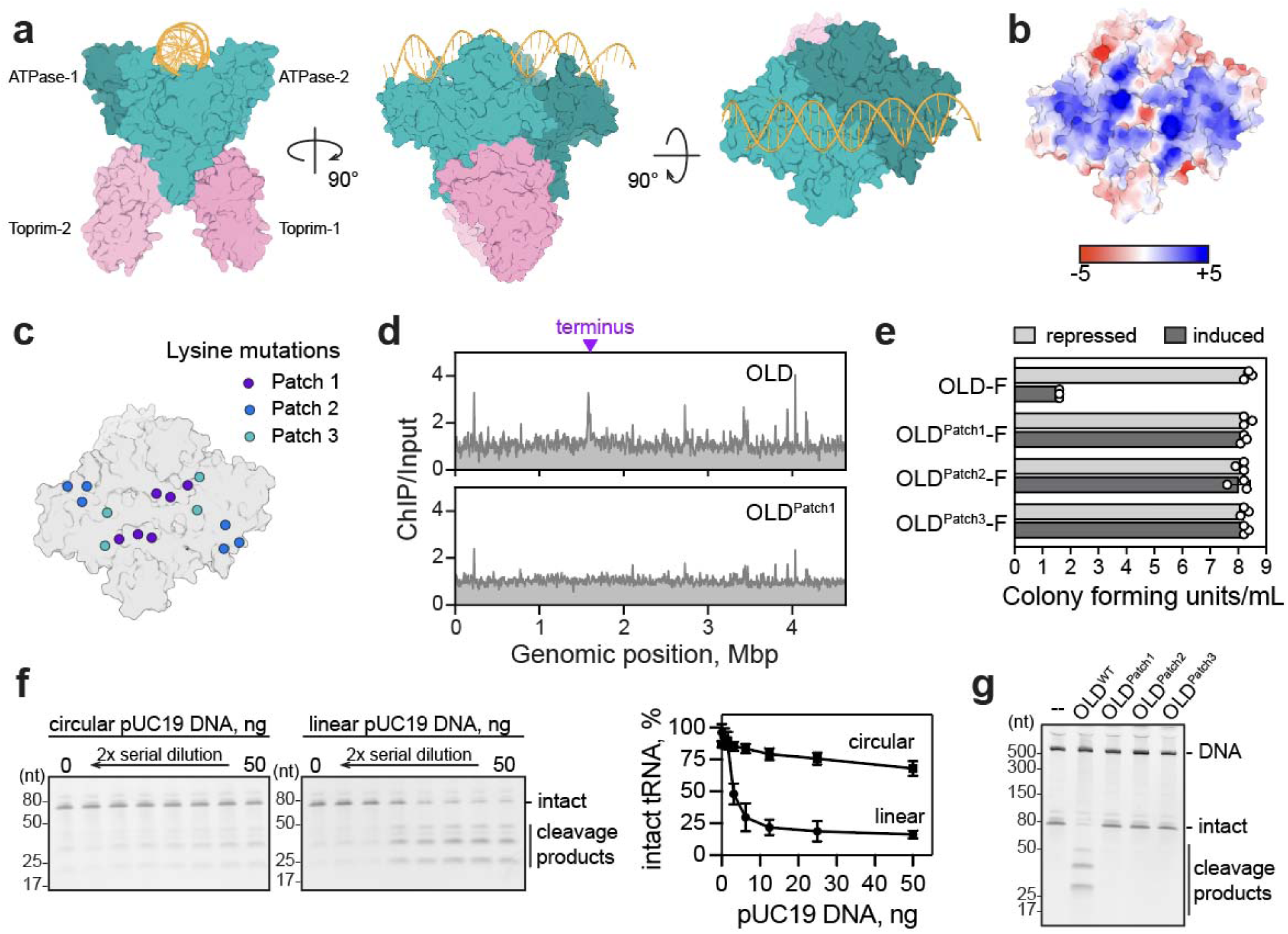
DNA-end recognition allosterically activates OLD’s tRNA nuclease activity. **(a)** Protenix structural model of an OLD dimer bound to ATP and a DNA duplex (yellow) [ptm: 0.6251, iptm: 0.5663]. The ATPase and Toprim domains of each subunit are labeled teal and pink, respectively. Subunit 1 is differentiated from Subunit 2 by a darker shade. **(b)** Electrostatic surface charge distribution of the dimerized OLD ATPase domains (*top view*). **(c)** Mutated lysine side chains (spheres) are mapped onto the basic channel formed by the dimerized OLD ATPase domains and color-coded by mutants. **(d)** Genome-wide ChIP-seq profiles of indicated FLAG-tagged OLD variants 40 minutes post-Gam induction. ChIP-seq was conducted as described in Fig. 2a, b. Representative of n = 2 biological replicates. **(e)** Toxicity quantification by spot dilution assay. Tenfold serial dilutions of wild-type cells harboring expression plasmids producing Gam with or without the indicated OLD proteins. Dilutions were spotted on LB agar with glucose or arabinose to repress or induce expression of FLAG-tagged proteins, respectively. Data, from three independent experiments, are the colony forming units per mL expressed as Log_10_. Plates were incubated at 42°C to induce Gam. **(f)** In vitro tRNA^ThrU^ cleavage by purified OLD in the presence of 0.5 mM ATP and 250 mM K^+^ across serial dilutions of circular or BamHI-HF-linearized pUC19 plasmid. **(g)** In vitro tRNA^ThrU^ cleavage by the indicated purified OLD variants in the presence of BamHI-HF-linearized pUC19 and ATP, as in panel **f**. In vitro experiments are representative biological triplicates.

To directly test this interpretation, we assessed how various nucleic acid substrates affect the ATP-driven activity of purified OLD towards its preferred tRNA^ThrU^ substrate^27^. At a near physiological concentration of 250 mM potassium, where basal tRNA cleavage is minimal, the addition of linearized plasmid DNA potently stimulated tRNA^ThrU^ cleavage (Fig. 3f and Fig. S4a, b). In contrast, circular plasmid DNA had minimal effects, and other nucleic acids—including single-stranded DNA, also a RecBCD substrate—failed to stimulate cleavage (Fig. S4c). Thus, specific activation of OLD by dsDNA ends can be recapitulated in vitro using purified components. Consistent with their proposed defect in DNA-end recognition, purified OLD^Patch1^, OLD^Patch2^ and OLD^Patch3^ all failed to cleave tRNA^ThrU^ in the presence of linear DNA (Fig. 3g). Mass photometry showed that the mutant proteins form dimers, ruling out global destabilization (Fig. S4d). However, these mutants did not assemble into higher-order oligomers observed for a small fraction of the wild-type enzyme (Fig. S4d), suggesting that DNA-end detection by the ATPase domain modulates oligomerization to allosterically activate the Toprim domain for tRNA cleavage.

### Individual DNA ends activate many OLD molecules at once

Individual peaks in our ChIP-seq data typically cover 5-7 kbp (Fig. 2d), an unusually large window for a protein that recognizes DNA ends. Although population-level heterogeneity in the location of DNA breaks could explain this wide distribution, the intrinsic dynamics of OLD on the chromosome must also play a role, as peak shape is noticeably altered by ATPase activity (Fig. 2d and Fig. S2d). Alternatively, this broad enrichment may reflect the simultaneous loading of multiple OLD complexes onto individual DNA molecules. To address these questions, we constructed *E.* coli expressing a functional OLD-mVenus fusion (Fig. S5a) and imaged these cells following Gam induction. While the OLD-mVenus signal was initially diffuse throughout the cytoplasm, it progressively reorganized into distinct foci upon Gam-mediated RecBCD inhibition (Fig. 4a,b). In contrast, OLD-mVenus remained diffuse in cells expressing a control peptide (Fig. S5b). Focus assembly persisted when cell division was blocked by cephalexin, demonstrating that they form independently of division-induced chromosome breakage (Fig. S5c).

**Figure 4.**
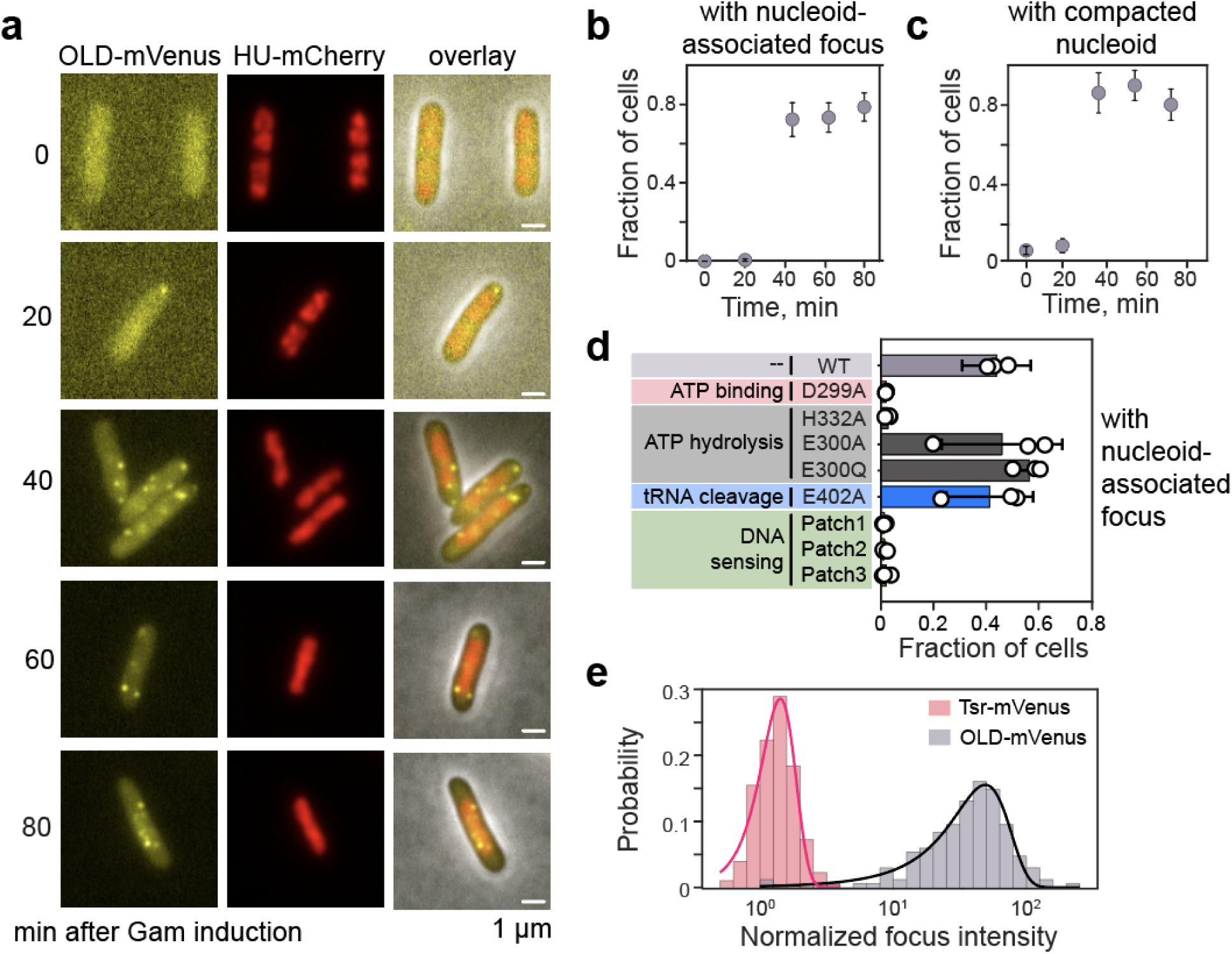
ATP-bound OLD molecules form foci upon Gam induction. **(a)** Representative images of cells expressing OLD-mVenus and HU-mCherry at indicated times after Gam induction. These images are overlaid with phase contrast images on the third column. **(b-c)** Quantification of images shown in (**a**). **(b)** Fraction of cells containing at least one nucleoid-associated OLD-mVenus focus (defined as located within or immediately adjacent to HU-mCherry signal). **(c)** Fraction of cells with a compacted nucleoid (defined as a nucleoid-to-cell area (NC) ratio < 0.45). Results are shown as the mean ± std across triplicates. Total n = 602, 511, 720, 569, and 686 cells were analyzed at t = 0, 20, 40, 60, and 80 min, respectively. **(d)** Fraction of cells containing at least one OLD-mStayGold focus for wild-type OLD and OLD mutants, measured 60 min after Gam induction. Results are shown as the mean ± std across triplicates, including n = 1340 (WT), 1149 (D229A), 1261 (H332A), 1077 (E300A), 921 (E300Q), 1840 (E402A), 909 (Patch1), 1015 (Patch2), and 1024 cells (Patch3). **(e)** Normalized focus intensities of single mVenus molecules determined from Tsr-mVenus photobleaching steps (gray, n = 269 foci) and individual OLD-mVenus foci (pink). Total n = 179 OLD-mVenus foci from 104 cells imaged at t = 60 and 80 min after Gam induction were analyzed. The solid line represents a Gaussian fit in log scale (µ = 1.1 for Tsr-mVenus and µ = 35 for OLD-mVenus).

The majority of OLD foci appear to associate with the nucleoid (Fig. 4a) and their appearance tracked the kinetics of nucleoid compaction (Fig. 4a-c)—a cytological indicator of global translational arrest in *E. coli^39^*—suggesting a functional link to DNA-end triggered activation of OLD’s tRNA nuclease activity (Fig. 3). To ask whether these foci correspond to the terminus-bound OLD observed by ChIP-seq, we investigated the biochemical requirement for focus formation. We reasoned that if these phenomena reflect the same underlying complex, they should share identical functional requirements.

We addressed this proposal by imaging nontoxic derivatives of an OLD-mStayGold fusion (Fig. S5a and Fig. S6a-c), which behaved similarly to OLD-mVenus (Fig. 4a-c). Consistent with our prior ChIP-seq results for terminus association, mutations that disrupt ATP sensing (H332A) or DNA binding abolished the formation of nucleoid-associated foci, whereas those that selectively inactivate ATP hydrolysis (E300Q/A) or tRNA cleavage (E402A) had no effect (Fig. 4d and Fig. S6d). Occasionally, we observed polar foci in mutants that failed to assemble on the nucleoid; the relevance of these foci is unknown. The perfect correlation with our ChIP-seq results indicates that the nucleoid-associated foci likely reflect terminus-bound OLD complexes. Moreover, a D299A substitution in the OLD Walker B motif—predicted to disrupt metal and ATP binding—abolished focus assembly in this assay (Fig. 4d). This result corroborates our previous interpretation that ATP binding, and the accompanying conformational change (Fig. 2d), are required for OLD to detect the DNA trigger at the chromosomal terminus.

Having established the functional relationship between the fluorescent foci and terminus associated OLD, we next quantified the copy number of OLD molecules in each focus. To do so, we used OLD-mVenus and calibrated their fluorescence signals against single molecules of Tsr-mVenus (Fig. S7a–e). We chose Tsr-mVenus as a standard because its localization within the chemoreceptor arrays renders it immobile during imaging. Mean focus intensity of OLD-mVenus was 42-fold higher than single Tsr-mVenus molecules (Fig. 4e), corresponding to roughly 21 OLD dimers per focus. Assuming each focus represents a single DNA end and each dimer occupies a 30-bp footprint per dimer (Fig. 3a), 21 dimers would occupy just ∼630 bp—far short of the ∼5,000-bp binding zone observed by ChIP-seq (Fig. 2c,d). Consequently, OLD is unlikely to coat the entire region as a contiguous polymer. This broad distribution could be explained by two alternative models: it may reflect a combination of limited polymerization and significant population-level heterogeneity in dsDNA endpoints, or it could arise from the dynamic loading of multiple independent OLD molecules through the DNA end. Because inactivating ATPase hydrolysis alone shifts the peak relative to the DNA end, we favor the latter model (See Discussion). Nonetheless, the capacity of multiple OLD to associate with a single DNA end may provide a built-in signal amplifier, allowing even a few DNA ends to activate a large number of OLD for rapid tRNA cleavage.

### Novel defense systems that trigger toxicity without RecBCD

To investigate whether the autoimmune risk posed by accumulated self-DNA extends broadly across bacterial defense mechanisms, we sought to identify additional systems that are influenced by RecBCD. We transformed a pooled plasmid library expressing genes from phage λ in MG1655 containing GFP or antiphage systems known to defend against λ for toxicity upon Gam-mediated RecBCD inhibition^40^. Plasmid depletion analysis revealed that Gam was specifically depleted in the presence of *E. coli* ECOR12 DRT3, Sir2+HerA, DUF4297+HerA, and Ppl systems, but not in MG1655 producing GFP, suggesting that these systems may become activated upon perturbation of RecBCD (Supplementary Table 1). Transformation assays with a single plasmid confirmed Gam is toxic to bacteria expressing these systems (Fig. 5a, b) Further testing revealed that Sir2+HerA likely senses Gam directly rather than RecBCD inactivation, as no toxicity was observed when RecBCD was inhibited by expressing T7 Gp5.9 (a DNA mimic like Gam) (Fig. 5a,b) or inactivated genetically (Fig. 5c, d). Of the remaining three systems that demonstrated toxicity upon T7 Gp5.9 expression—marked by decreased colony formation—only DRT3 and Ppl showed toxicity in a temperature-sensitive *recBC* mutant (Fig. 5c, d). Our finding is consistent with recent work demonstrating DRT3’s toxicity in *recBCD* mutants and the proposal that RecBCD suppresses toxicity by degrading DNA duplex synthesized by DRT3^41^. In contrast, Ppl likely induces toxicity by attacking self-DNA ends at the replication terminus as it does during infection^42^. Notably, Ppl harbors a Rad50-like ATPase domain similar to those of OLD and Gabija, both of which induce toxicity in RecBCD-deficient bacteria^16,20,24^.

**Figure 5.**
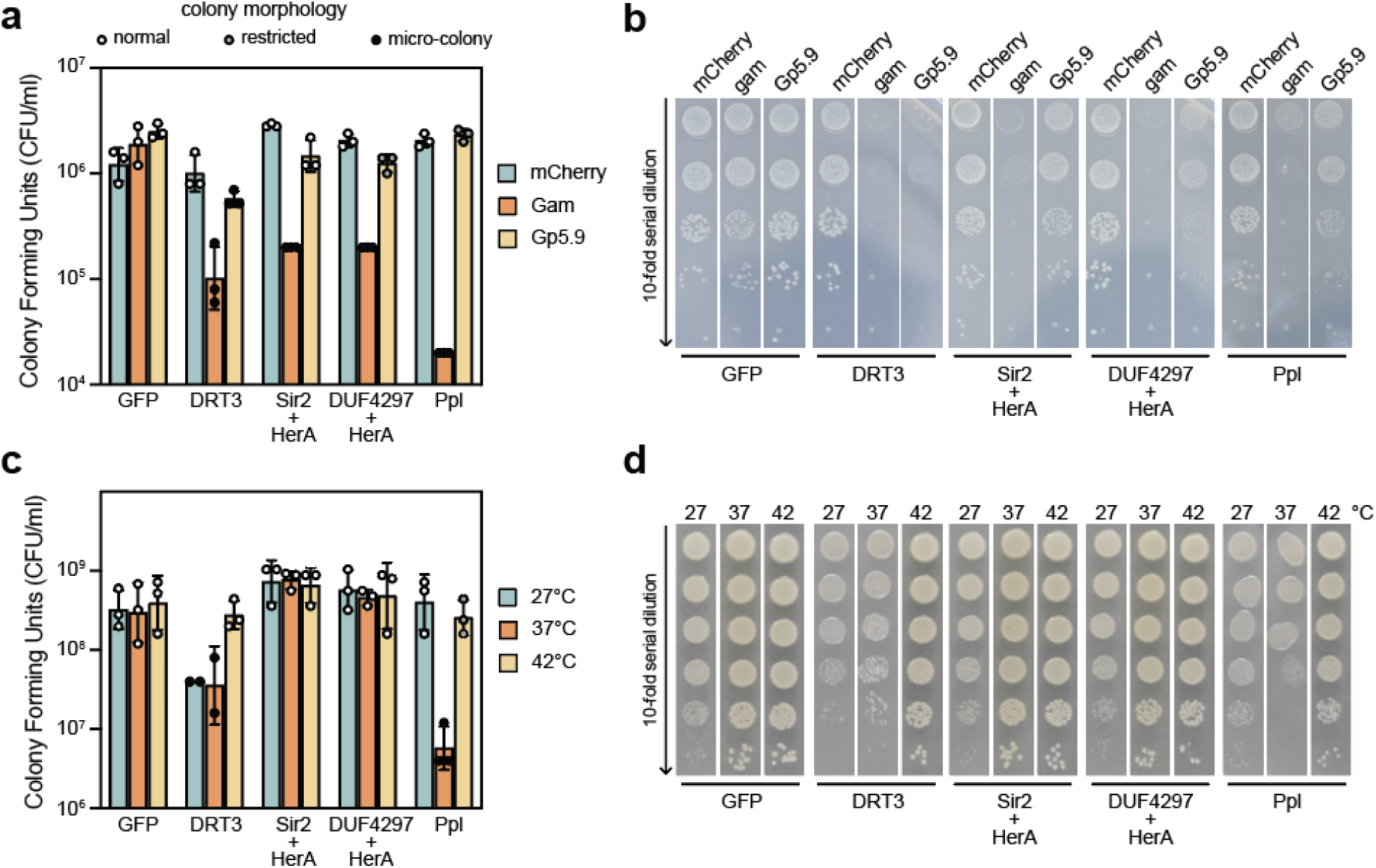
Identification of new immune systems that are toxic without RecBCD. (**a**) Quantification of colony forming units (CFU) of MG1655 expressing GFP or indicated defense system recovered after transformation with a plasmid expressing mCherry, λGam, or T7 Gp5.9, and cultivated on medium containing Carb100 and 50 µM IPTG. Colony morphology was categorized by colors: normal (white circles), restricted growth characterized by reduced colony size (gray circles), and micro-colonies exhibiting pin-prick size and/or transparent appearance (black circles). (**b**) Representative images of recovered CFU. **(c**) Viability of *E. coli* strain SK128 harboring temperature-sensitive *recB270 and recC271* alleles producing the indicated defense systems or the GFP control at 27°C, 37°C or 42°C. Freshly transformed colonies were grown in LB with Carb100 at 27°C. OD-normalized cultures were then serially diluted ten-fold and spotted on LB agar containing Carb100 and 0.2% glucose and incubated at the indicated temperatures. Colony morphology was scored as described in (**a**). Representative images of CFU are shown in (**d**). Data are the mean ± standard error of the mean (SEM) triplicates.

## Discussion

Although sensing of genome integrity has emerged as a recurring theme in bacterial immunity, known systems are thought to respond to rampant DNA damage or unique intermediates generated during phage replication^13–15,43–45^. Here, we define physiological DNA ends as a distinct class of immune trigger. These DNA ends, which are routinely generated during replication termination, normally evade immune detection because they are rapidly cleared by the housekeeping RecBCD helicase-nuclease. In its absence, they persist and inappropriately activate the OLD antiphage defense, driving autoimmune toxicity. This finding reveals a unifying principle across kingdoms: essential cellular processes generate immunogenic byproducts that must be actively neutralized to prevent self-targeting. Just as the TREX1 exonuclease and DBR1 debranching enzyme clear immunostimulatory cytosolic DNA and plant intronic lariats^9–12,46^, RecBCD acts as an autoimmune safeguard in bacteria, resolving the inherent molecular conflict between normal physiology and antiviral defense.

Replication termination frequently generates linear dsDNA with exposed ends^37,47^, and their processing by RecBCD is essential for sister chromosome segregation^34,48^. Without it, chromosomes remain linked, causing severe terminus DNA loss during cell division^33,35^. By synchronizing RecBCD inhibition, we showed that OLD detects the initial uncleared dsDNA ends, rather than the ensuing DNA damage. Supporting this, OLD robustly recognizes ectopically induced site-specific dsDNA breaks and forms foci even when cell division—and thus DNA loss—is blocked. Importantly, genetically abolishing DNA-end sensing prevents OLD activation, providing, to our knowledge, the first direct evidence of autoimmune activation by physiological host DNA ends.

The immune safeguard function we describe here adds a new dimension to host nucleases’ established roles in bacterial immunity. While nucleases like ExoI and RecBCD separately prevent the autoactivation of distinct defense-associated reverse transcriptases, they do so by clearing the nucleic acids synthesized by the defense systems themselves^41,49^. Beyond these immune safeguard functions, host nucleases—most prominently RecBCD—are harnessed by immune systems like CRISPR and prokaryotic Argonaute for memory acquisition^50–53^ and single-stranded DNA guide generation^54^, respectively. Together, these findings highlight the frequent co-option of host nucleases for immune function and the multifaceted roles of RecBCD in bacterial immunity.

Our data indicate that OLD detects dsDNA ends using a channel formed across its dimerized ABC-type ATPase subunits, analogous to how the eukaryotic Rad50 recognizes its target (Fig. 3). This conserved mechanism of DNA-end sensing is likely pervasive, as bacterial defense systems with ABC-type ATPases are widespread^24,44,55,56^. Indeed, these diverse systems share mechanistic parallels. Like OLD, Lamassu is allosterically activated by dsDNA ends in vitro^43–45^. In cells lacking RecBCD, the OLD, Gabija^20^ and Ppl (as we show here) systems all trigger toxicity (Fig. 5). Furthermore, during infection, phage can bypass both OLD and Ppl defenses by inactivating their own homing endonucleases, which presumably reduces the generation of immunogenic DNA ends^42,57^. Finally, OLD, Lamassu, and Gabija localize to the replication origins of certain phages^20,44,57^, implicating recruitment by stem-loops or DNA ends formed during viral replication initiation. A critical future direction will be to investigate whether these diverse systems universally depend on RecBCD, or if they have evolved distinct regulatory mechanisms to prevent aberrant activation by termination-derived self-DNA ends.

*E. coli* cells harbor at most four complete chromosomes^33^, limiting the number of DNA ends that can form. Given this, the severe OLD-mediated autoimmune response of *recBCD* mutants is surprising^16,27^ (Fig. 1). Our work reveals that dozens of OLD molecules can engage a single DNA end at once, providing a potential signal amplification mechanism that explains this sensitivity (Fig. 4). Physiologically, this amplification explains how OLD rapidly aborts λ infection minutes after entry, well before significant viral replication occurs^22,58,59^. While the precise mechanism of this multivalent association requires further investigation, our mutagenesis data (Fig. 2d and 4d) support a dynamic loading model (Fig. 6): ATP binding licenses recognition and loading at a dsDNA end. Blocking ATP hydrolysis (OLD^E300Q^) shifts the ChIP-seq binding profile further from the ends (Fig. 2d and S2d), suggesting that loaded complexes diffuse away from the dsDNA end until ATP hydrolysis triggers their dissociation. This clears the DNA end for loading more OLD molecules, leading to multivalent association. In this model, the relative rates of one-dimensional diffusion and ATP hydrolysis dictate the length of DNA occupied by OLD while the relative rates of loading and ATPase hydrolysis dictate the number of OLD molecules per DNA end. Notably, because ATP-hydrolysis-negative OLD^E300Q^ is nontoxic (Fig. S6a), DNA-binding itself does not unleash tRNA nuclease activity. Whether OLD cleaves tRNA while transiently staying on the DNA after ATP hydrolysis, or whether this cleavage occurs off-DNA, remains to be determined.

**Figure 6.**
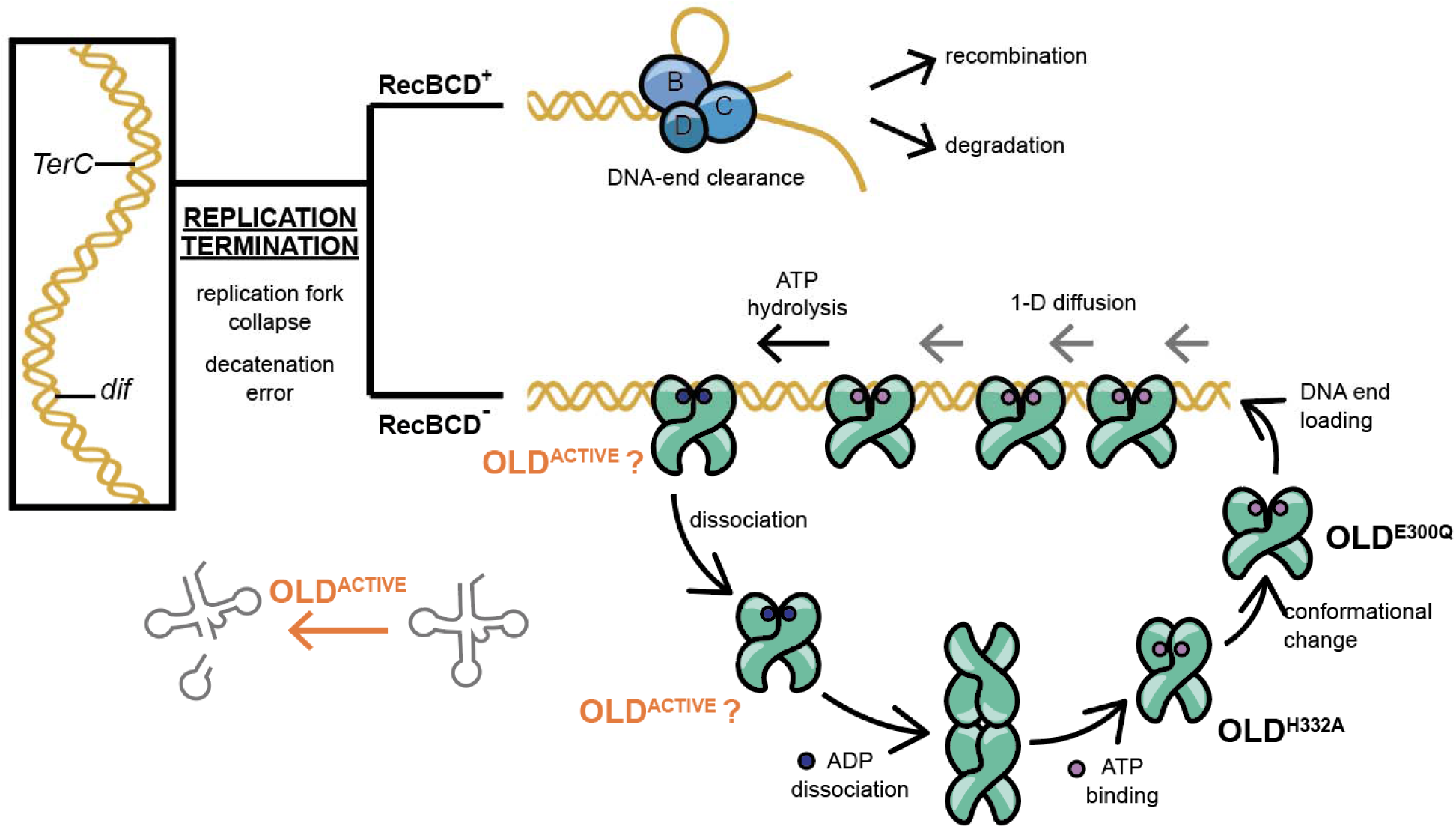
Model of OLD-mediated toxicity in the absence of RecBCD. Schematic showing the fate of DNA ends generated during replication termination in wild-type and a *recBCD* mutant. RecBCD eliminates these DNA ends by initiating homologous recombination or through degradation. A hypothetical model depicts how the ATPase cycle of OLD controls multivalent association with persisting DNA ends through a dynamic end-loading mechanism. OLD variants that trap specific intermediate states within the reaction cycle are indicated. An inactive tetrameric state, reported in a parallel study, is also depicted.

Discovered in 1969 as a toxic factor that prevents P2 lysogenization of *recBC* mutant hosts^16^, the OLD antiphage system has long presented a mysterious case of bacterial autoimmunity. Because RecBCD is central to DNA repair^29^, and OLD sensitizes cells to X-ray it was thought that this toxicity arose from the cell’s inability to contain OLD’s proposed DNA exonuclease function^25,26^. We recently changed this view, demonstrating that OLD is actually a tRNA nuclease that halts translation by amputating the anticodon stem of specific host tRNAs^27^. Here, together with a parallel we bring this story full circle by uncovering the endogenous trigger signal in *recBCD* mutants. More broadly, resolving this age-old puzzle also reveals immunological principles conserved across prokaryotes and eukaryotes.

The molecular basis of OLD toxicity in *recBCD* mutants has evaded understanding for over fifty years. Last year, we provided a piece of this puzzle, establishing that OLD functions as a tRNA nuclease that halts translation by amputating the anticodon stem of host tRNAs^27^. Moreover, by showing that this cell death is suppressible by resistant phage tRNAs^27^, we established the mysterious tRNA inactivation observed decades ago as the biochemical basis of toxicity^22,58^. Here, alongside a parallel study^28^, we complete this model by uncovering the endogenous trigger signal: the accumulation of DNA ends in *recBCD* mutants. This end-dependent activation elegantly accounts for the long-standing observation of X-ray sensitivity in bacteria encoding OLD^60^. More broadly, resolving this classic puzzle unveils deeply conserved immunological principles across prokaryotes and eukaryotes.

## Methods

### General methods for cell-based assays

*E. coli* strains and bacteriophage used in this manuscript are summarized in Supplementary Table 2, respectively. Sequences of plasmids used in this manuscript are listed in Supplementary Table 3. Plasmids were ordered from GenScript or constructed in-house using NEBuilder HiFi DNA Assembly mix (NEB). DNA fragments were amplified with Q5 High-Fidelity Polymerase (NEB) or purchased as gBlocks (IDT). Unless mentioned otherwise, all bacterial strains used in cell-based assays are derivatives of wild-type *E. coli* strain MG1655. Bacterial strains were cultured in lysogeny broth (LB-Miller; Teknova). Antibiotics were added to the growth media as needed at the following concentrations: 50 µg ml^-1^ kanamycin (Kan50), 100 µg ml^-1^ carbenicillin (Carb100), 100 µg ml^-1^ ampicillin (Amp100), 15 or 20 µg ml^-1^ chloramphenicol (Cm15 or Cm20) or 10 µg ml^-1^ tetracycline (Tet10). Expression from the *P_BAD_* promoter was repressed with 0.2% glucose and induced with 0.2% arabinose. For genes under the control of temperature-sensitive *P_cI857_* promoter, cultures were grown at 30°C for repression and shifted to 37°C or 42°C (as indicated) to induce expression.

### Spot dilution assays

Several freshly transformed colonies were used to inoculate 3 ml of LB supplemented with the appropriate antibiotics and glucose and grown at the indicated temperatures until OD_600nm_= 0.5-1.5. Following dilution to OD_600nm_= 0.5, the culture was serially diluted 10-fold in LB. 5 µl of each dilution was spotted on LB agar supplemented with antibiotics and either glucose or arabinose when appropriate. The plates were then incubated overnight at the indicated temperature before images were recorded. Images of the agar plates were taken with a Nikon D3500 camera with a Nikon AF-S DX Micro NIKKOR 40mm f/2.8G Lens.

### Protein expression and purification

Freshly transformed One Shot BL21 Star (DE3) (Invitrogen) colonies harboring pNEB265 (His6-MBP-SUMO-OLD-TwinStrepTag), pNEB486 (His6-MBP-SUMO-OLD^Patch1^-TwinStrepTag) and pNEB487 (His6-MBP-SUMO-OLD^Patch2^-TwinStrepTag) or pNEB488 (His6-MBP-SUMO-OLD^Patch3^-TwinStrepTag) were scraped and used to inoculate 1 L of LB (Teknova) supplemented with Kan50 at OD_600nm_ of 0.05. The culture was grown at 37°C with shaking (200 rpm) until OD_600nm_ ∼ 1. The culture was chilled to 4°C and the protein was induced with 0.5 mM IPTG (Teknova) and grown overnight at 18°C with shaking at 200 rpm. The cells were harvested by centrifugation at 4000 x g for 20 min at 4°C and stored at −80°C. The cell pellet was resuspended in 40 ml Lysis buffer (20 mM Tris HCl buffer pH 8.0, 250 mM NaCl, 10 % glycerol, 1 mM DTT) that was supplemented with 1 tablet of EDTA-free complete Protease Inhibitor Cocktail (Roche). Cells were lysed by sonication using an Qsonica Q700 sonicator (50% power, 3 sec ON and 6 sec Off for a total of 2 min). Cell debris was removed by centrifugation at 13,000 x g for 30 min and the clarified lysate was passed over 3 ml Streptactin-XT resin (IBA Lifesciences) pre-equilibrated with lysis buffer by gravity flow. The resin was washed with 5 column volumes of lysis buffer supplemented with 1 M NaCl and protein was eluted using lysis buffer supplemented with 50 mM biotin. Protein was concentrated using Amicon Ultra Centrifugal Filter, 30 kDa MWCO (Millipore UFC9030), and purified using a Superdex16/600 S200pg column (Cytiva) in 20 mM Tris-HCl pH 8, 1 mM NaCl, 10% glycerol, and 1 mM TCEP. Fractions containing OLD and its mutant forms were pooled, dialyzed into storage buffer (20 mM Tris-HCl pH 8, 200 mM NaCl, 10% glycerol, and 1 mM TCEP), concentrated, aliquoted and flash frozen.

### tRNA^ThrU^ cleavage assay

Cleavage assay was performed with the *E. coli* tRNA^ThrU^ substrate, which was prepared by in vitro transcription using the HiScribe® T7 Quick High Yield RNA Synthesis Kit (NEB) as described previously^1^. Following purification using the Monarch RNA clean up kit (NEB), tRNA^ThrU^ was diluted into 20 mM Tris, 10 mM Mg-acetate, and 50 mM K-acetate buffer pH 7.9 to a final concentration of 80 ng/µl. To ensure folding, diluted tRNA was incubated at 95 °C for 5 min and allowed to cool slowly to room temperature prior to use in the cleavage assay.

Circular pUC19 plasmids (NEB) were treated with T5 exonuclease (NEB) in CutSmart buffer to remove residual linear and nicked DNA. Linear pUC19 plasmids with different ends were prepared by treatment with 20 U of BamHI-HF (NEB), 20 U of SmaI (NEB), or 20 U of PstI-HF (NEB) in CutSmart buffer. Reactions were incubated at 37 °C for 1 h, subsequently cleaned up using Monarch Spin PCR & DNA Cleanup Kit (NEB) and eluted in 50 µL of water. For alternative nucleic acid substrates, M13mp18 single-stranded DNA (NEB) was diluted to 50 ng/µl, *egfp* mRNA (from Dillon Nye) was diluted to 50 ng/µl, and tRNA mimic (tRNA like structure of the Turnip Yellow Mosaic Virus) and viral dsRNA (adenovirus virus-associated RNA1) were both diluted to 50 ng/µl. See Supplementary Table 4 for sequences of these RNAs.

In 10 µl reactions, 80 ng tRNA^ThrU^ was mixed with 950 nM OLD in Reaction buffer (20 mM Tris-HCl, 10 mM Mg-acetate, 250 mM K-acetate, pH 7.9) supplemented with 0.5 mM Mg-ATP. When indicated, RNA and DNA co-factors were added. For time course experiments, this reaction was scaled up proportionally. Reactions were incubated at 37°C for 1 hour, quenched by adding 25 mM EDTA supplemented with 6 U of Proteinase K (NEB), diluted two-fold with 2x TBE-urea sample buffer (Thermo). 4 µL of reaction was run on a 15-well 15% TBE-Urea gel at 180 V for 45 minutes. Gels were stained with SYBR gold (Thermo) in 1xTBE buffer (Invitrogen) before imaging using an Amersham Typhoon in fluorescence mode with the Cy2 filter.

Cleavage of tRNA^ThrU^ was quantified by analyzing gel images using the open-source software Fiji^2^. The fraction of intact tRNA^ThrU^ was quantified by dividing the band intensity over the intensity of the entire lane following background subtraction.

### Mass photometry analysis

Purified OLD proteins were diluted to 72 nM in Reaction buffer supplemented with 1 mM Mg-ATP. Mass photometer (Refeyn 2MP) was turned on an hour before use. 10 µL of buffer was spotted on slides to calibrate the focus before mixing with 10 µL of protein in buffer for a final concentration of 36 nM. Videos were recorded for 60 sec, and protein counts were graphed and analyzed using the Refeyn Mass photometer analysis software.

### Plaque Assay

Freshly transformed *E. coli* colonies were inoculated into 2 mL of LB with arabinose, Cm and Tet and grown at 37°C with shaking until OD_600nm_ > 1. 100 µL of the culture was mixed with 4 ml of 0.75% soft agar in LB with 0.2% arabinose, 5 mM Mg-OAc and 5 mM CaCl_2_. The mixture was poured onto pre-warmed LB plates; after 20 min, 2 µL of ten-fold serially diluted phages lysates of λ*vir* were spotted. The plates were incubated overnight at 37°C.

### ChIP-seq analysis

Freshly transformed MG1655 colonies harboring pNEB64 (encoding temperature-inducible Gam) and plasmids encoding FLAG-tagged OLD variants or GFP were scraped, suspended in LB, and inoculated into 50 mL LB containing Cm15 and Tet10. Cultures were incubated at 30°C with shaking (200 rpm). At OD_600nm_ = 0.2, arabinose was added to induce OLD-F production, and growth continued to OD_600nm_ = ∼1.0. A 20 mL aliquot of the culture was collected as the time-zero sample. The remaining culture was diluted two-fold into pre-warmed (42°C) LB supplemented with Cm, Tet and arabinose (OD_600nm_ = 0.5) and incubated at 42°C with shaking for the indicated times. For site-specific DSB induction using I-*Sce*I, *E. coli* strains HCL252 (genomically encoding an ectopic *sce* cleavage site and *P_BAD_*-driven I-*Sce*I) or HCL254 (encoding the *sce* site alone) were transformed with pNEB502 (constitutively expressing OLD^E402A^-F) and pNEB64 (temperature-inducible Gam). Cells were grown as described above, except arabinose was added at OD_600nm_ = 1 (immediately before dilution and the temperate upshift to 42°C) to simultaneously induce I-*Sce*I and Gam. Cultures were harvested after 1 hour.

We cross-linked 20 mL of the culture by rotating with 1.8 mL of formaldehyde (36.5-38% in H2O; Sigma) in a 50 ml conical tube at room temperature for 30 min. We then quenched the reaction by adding 1 mL of 2.5 M glycine and rotating for 5 min. Finally, we harvested cells by centrifugation (5000 x *g*, 10 min, 4°C), washed them three times with 15 mL ice-cold PBS (Thermofisher), flash frozen on dry ice, and stored them at −80°C.

### ChIP

To lyse the cells, we resuspended the cell pellet in 400 µL of buffer (12.5 mM Tris-HCl pH 8, 12.5 mM EDTA pH 8, 62.5 mM NaCl, 25% sucrose) supplemented with 100 µL of lysozyme (25 mg/ml) and incubated at 37°C with shaking at 300 rpm for 30 min. We then added 500 µL of 2x IP buffer (100 mM Tris pH 7, 10 mM EDTA, 300 mM NaCl, 2% Triton X-100) and completed lysis via sonication using a Qsonica Q700 sonicator (30% power, 5 second ON, 15 second OFF for 2 min total ON-time). After centrifugation (23,000 x *g*, 20 min, 4°C), the supernatant was recovered, combined with 50 µL of Protein-G Magnetic Beads (NEB; pre-equilibrated in 1x IP buffer), and incubated at 4°C with gentle rotation for 1.5 hours. 20 µL of the pre-cleared lysate was combined with 200 µL of elution buffer (50 mM Tris-HCl pH 8, 10 mM EDTA, 1% SDS) and set aside as the Input sample. The remaining lysate was incubated with 20 µL of ANTI-FLAG M2 monoclonal antibody (Sigma, F1804) at 4°C with rotation overnight. Immunoprecipitated complexes were captured by adding 50 µL of Protein-G Magnetic Beads in 1x IP buffer and rotating at 4°C for 1.5 hours. Beads were collected magnetically and washed three times with 1 mL of 1xIP buffer, followed by three washes with 1 mL Wash Buffer (50 mM Tris-HCl pH 7.5, 1 mM EDTA, 500 mM NaCl, 1% Triton X-100). We performed a final wash with 1 mL of TE buffer (5 min rotation at room temperature) and eluted the ChIP DNA by incubating the beads in 250 µL of elution buffer (50 mM Tris-HCl pH 8, 10 mM EDTA, 1% SDS) for 10 min at 65°C. After magnetic separation, we incubated the ChIP and Input samples overnight at 65°C to reverse cross-linking.

On the following day, sample volumes were adjusted to 500 µL with TE buffer pH 8. Samples were treated with 2.5 µl RNAseA (0.2 mg/mL final; NEB) at 37°C for 30 min, followed by treatment with 5 µL of Proteinase K (0.2 mg/mL final; NEB) at 37°C for 30 min. Phenol:Chloroform:Isoamyl Alcohol (25:24:1; 500 µL) was added, vortexed, and centrifuged at 13,000 x *g* for 5 min. The aqueous phase was transferred to a fresh tube and mixed with 50 µL of 3 M sodium acetate and 1 µL of glycogen (20 mg/mL). Samples were vortexed, mixed with 1 mL 100% ethanol, vortexed again, and incubated at −80°C for 30 min. The samples were centrifuged at 4°C for 30 min at 15,000 x *g*. The supernatant was discarded, and the DNA pellets were washed twice with 1 mL of 70% ethanol. Air-dried pellets were resuspended in 100 µL nuclease-free water (NEB). DNA was purified using 1.6x (160 uL) NEBNext® Sample Purification Beads, and eluted in 53 µL of nuclease-free water. Input DNA (1 µL diluted with 49 µL of water) and 50 µL of immune-precipitated DNA were subjected to end repair and dA-tailling using NEBNext End Prep, by adding 7 µL of 10x End Repair Reaction buffer and 3 µL of End Prep Enzyme mix, followed by two 30-min incubation at 20°C and 65°C, sequentially.

NEBnext Adaptor for Illumina was diluted 1:5 in 10 mM Tris-HCl pH 8, 10 mM NaCl and 2.5 µL of diluted adaptor was added directly into each end-prepped reaction, followed by a master mix of 30 µL NEBnext Ultra II ligation Master mix and 1 µL of NEBNext Ligation Enhancer, and incubated at 20°C for 15 min. 3 µL of USER enzyme (NEB) was added directly to each ligation reaction and incubated at 37°C for 30 min. Adaptor ligated DNA was cleaned up using 0.9x NEBNext® Sample Purification Beads to remove unligated adaptors and resuspended in 17 µL of nuclease-free water. 15 µL of DNA was mixed with 25 µL NEBNext Q5 Hot start HiFi 2xPCR master mix and 10 µL of NEBNext Illumina primers. PCR products were pooled and cleaned up using 0.8x NEBNext® Sample Purification Beads prior to sequencing with NovaSeq 6000 or Element Aviti.

### ChIP data processing

Raw ChIP-seq reads were trimmed using Cutadapt (version 4.5). Trimmed reads were mapped to *E. coli* MG1655 reference genome (NC_000913) or NC_000913 modified with *sce-*frt-*kan*-frt insertion at with STAR (version STAR_2.5.2b) with --alignIntronMax 1 argument to turn off splicing alignments. SAMtools software package (version 1.17) was used to select for primary alignments with a mapping quality > 10, and subsequently to sort and index-filter the mapped reads. Alignments were normalized by total read counts and converted into 1-bp-bin bigwig files using deepTools2 command bamCoverage. Unless indicated otherwise, ChIP coverage was normalized to Input coverage using custom Python scripts. The genomic region encoding *araC* (positions 70,000–71,300) was masked because *araC* is also encoded on a plasmid used in these experiments. For experiments involving I-SceI cleavage, genomic regions spanning positions 1,380,000–1,413,000 and 1,979,500–1,981,500 were also masked due to sequence discrepancies between the experimental *Escherichia coli* MG1655 strains and the reference genome. Genome-wide coverage plots are displayed in 300-bp bins, whereas zoomed views are displayed in 100-bp bins.

### Fluorescence microscopy

Cultures were grown overnight in LB supplemented with appropriate antibiotics and 0.2% glucose at 30°C with shaking (220 rpm). The following day, cultures were diluted 1:100 into fresh LB medium supplemented with appropriate antibiotics and 0.2% (w/v) arabinose to induce OLD expression. Expression of temperature-inducible constructs was initiated by shifting cultures from 30°C to 42°C, with the temperature shift designated as time zero. Samples were collected immediately before the temperature shift (t = 0) and every 20 min after. They were immediately imaged after being collected. For imaging, cells were immobilized on 1% (w/v) agarose pads prepared with LB and supplemented with the appropriate antibiotics and inducers. Pads were sealed beneath #1.5 coverslips using VALAP and imaged within 10 min of mounting. Fluorescence microscopy was performed on a Nikon Ti-2 microscope equipped with a Plan Apochromat objective 100×/1.45 NA oil-immersion phase objective, a SOLA SE II 365 LED illumination system (Lumencor), and an ORCA-R2 CCD camera (Hamamatsu). Images were acquired using Nikon NIS-Elements software. mCherry fluorescence was used to visualize the nucleoid, while mVenus or mStayGold fluorescence was used to visualize OLD.

### Cephalexin treatment

For inhibition of cell division, cephalexin (5 mg/mL in DMSO) was added immediately before imaging to a final concentration of 50 μg/mL. Cells were prepared and imaged identically to concurrent untreated samples.

### Image analysis and nucleoid-to-cell ratio measurements

Cell contours and nucleoid boundaries were determined using *Oufti* as previously described^3^. The same segmentation parameters were applied across all datasets. Cell area and nucleoid area were extracted from *Oufti*, and the nucleoid-to-cell area (NC) ratio was calculated as NC ratio = nucleoid area / cell area as previously described^4^.

### Classification of OLD localization

Fluorescent foci were manually quantified using a custom MATLAB graphical user interface (GUI). For each image, segmented cell meshes generated from *Oufti* were overlaid onto fluorescence or phase contrast images. Individual cells were displayed within the GUI and manually recorded with the cell number and localization of fluorescent foci. Cells were categorized into four groups based on focus localization: (i) no foci, defined as cells lacking detectable puncta; (ii) polar focus only, defined as cells containing only polar puncta located at the poles; (iii) cytoplasmic focus only, defined as cells containing only cytoplasmic puncta; and (iv) both, defined as cells containing both polar and cytoplasmic puncta. For each cell, the resulting foci classification was entered into the GUI using dedicated counters and automatically recorded in a results table. The GUI compiled classifications across all cells and generated histograms summarizing focus frequency and localization.

### mVenus single-molecule fluorescence quantification

Bright-field and fluorescence images were acquired using a Nikon Ti-2 microscope equipped with a 100x/1.49 NA TIRF objective (Nikon), a LUNF-XL laser box (Nikon), and an EMCCD camera (Andor iXon Ultra). Fluorescence images were acquired using 514-nm excitation at 1.38 mW/cm^2^ under epi-illumination, allowing fluorescence to be collected throughout the cell volume. A custom shutter was placed between the sample and the condenser to prevent reflected laser light from reaching the sample.

To determine the fluorescence intensity of a single mVenus molecule, *E. coli* cells expressing Tsr-mVenus (strain SK338) were grown at 37°C in MOPS EZ Rich Defined Medium (Teknova) to reduce cellular autofluorescence. Fluorescence images were acquired continuously for at least 1 min with an exposure time of 100 ms to capture single-step photobleaching events. Measured fluorescence intensities were converted to photon counts using the EMCCD camera specifications. The fluorescence intensity corresponding to a single mVenus molecule was determined from the photon numbers involved in single-step photobleaching events.

For OLD-mVenus measurements, *E. coli* cells (SK821) were grown in MOPS EZ Rich Defined Medium at 30°C and shifted to 42°C to induce Gam expression. Snapshot fluorescence images were acquired using the same imaging conditions used for Tsr–mVenus. Individual OLD-mVenus foci were manually identified using a custom MATLAB graphical user interface, and the integrated fluorescence intensity of each focus was calculated by summing the pixel intensities within a 5 × 5-pixel region centered on the focus. Local background fluorescence was estimated from three manually selected, signal-free regions within the same cell. The mean background intensity per pixel was multiplied by the area of the focus region and subtracted from the integrated focus intensity. The number of OLD-mVenus molecules in each focus was estimated by dividing the background-corrected focus intensity by the experimentally determined fluorescence intensity of a single mVenus molecule.

### Transformation efficiency assay

*E. coli* MG1655 expressing GFP, or DRT3, Sir2+HerA, DUF4297+HerA, or Gao_ppl were made electrocompetent by washing pelleted mid-log bacteria with ice cold water two times, and resuspending in ice cold 10% glycerol. 40 ng of control mCherry, λGam, or T7 Gp5.9 plasmid was electroporated into 50 μL of competent cells. Cells were then incubated in 1 mL of S.O.C. for 1 hour at 37 °C with shaking at 220 rpm, and subsequently serially-diluted and plated on MMCG agar (47.8 mM Na_2_HPO_4_, 22 mM KH_2_PO_4_, 18.7 mM NH_4_Cl, 8.6 mM NaCl, 22.2 mM Glucose, 2 mM MgSO_4_, 100 μM CaCl_2_, 3 μM Thiamine, Trace Metals at 0.1×, and Trace Metals Mixture T1001, Teknova, final medium concentration: 8.3 μM Ferric chloride, 2.7 μM Calcium chloride, 1.4 μM Manganese chloride, 1.8 μM Zinc Sulfate, 370 nM Cobalt chloride, 250 nM Cupric chloride, 350 nM Nickel chloride, 240 nM Sodium molybdate, 200 nM Sodium selenite, 200 nM Boric acid), supplemented with 20 μg/mL chloramphenicol, with or without 50 μM IPTG. Transformation efficiency was assessed by counting colony forming units after overnight incubation at 37 °C. In instances where single colonies were not distinguishable at the most dilute spot with visible bacteria, 10 colony forming units (CFU) were recorded. When no colonies were visible at the most concentrated spot, 0.9 CFU were recorded.

### Plasmid depletion assays

100 ng of pooled plasmids expressing every open reading frame (ORF) from phage λvir was transformed into *E. coli* MG1655 expressing GFP, or DRT3, Sir2+HerA, DUF4297+HerA, or Gao_ppl^5^. After one hour of recovery in S.O.C. at 37 °C and 220 rpm, cells were plated onto MMCG agar containing 50 μM IPTG and Cm20. For MG1655 expressing GFP, the transformation was also plated on MMCG agar containing 20 μg/mL Cm20 without IPTG. After 18 hours of growth at 37 °C, plasmids from surviving colonies with isolated by miniprep. 20 μL of miniprep was sequenced using 200Mbp Illumina Sequencing (SeqCenter). Reads were mapped to a λvir reference sequence using the *Map to Reference* feature of Geneious Prime after trimming reads using trim sequences: AGATGTGTATAAGAGACAG (adapter); ATAACAATTTCACACGCTAGCAGGAGGAATTCACC (5ʹ vector backbone); TGGCTCCAGCTTGCGGCCGCTTA (3ʹ vector backbone). Transcripts Per million (TPM) was used to compare relative read abundance between ORFs in different strains. ORF abundance was compared between uninduced MG1655 and MG1655 under IPTG induction. ORFs that had less than 10 reads under non-inducing conditions in MG1655 and those with 10-fold decrease in reads under IPTG induction were considered as growth inhibitory under induction and removed from further analysis. ORF abundance in MG1655 expressing defense systems was then compared to ORF abundance in uninduced MG1655 conditions.

## Data Availability

Data generated during the study is available in NCBI SRA (SUB16435554).

## Reagent and Code Availability

Reagents and code used during this study are available upon reasonable request to the corresponding authors [H.C.L. or S.K.].

## Acknowledgements

We thank the NEB leadership for their support of this project, Susan Rosenberg and for *E. coli* strains encoding I-SceI, NEB’s Sequencing Core led by Dora Posfai for help with high-throughput sequencing, Ian Rodney (Laub Lab, MIT) for λ*vir*, Matt Szucs for aliquots of the tRNA-like structure of the Turnip Yellow Mosaic Virus and adenovirus virus-associated RNA1, and Dillon Nye for *egfp* mRNA.

The work was supported by NIH (R35GM143203) and NSF Science and Technology Center for Quantitative Cell Biology (2243257) to S.K., Burroughs Wellcome Fund PATH Award (1186087) to A.T.W., and NIH (R35GM118120) to G.R.S.. New England Biolabs paid the salaries of A.A.G., I.U., M.S., C.T. and H.C.L. and funded their research.

## Author Contributions

A.A.G. purified proteins, performed in vitro transcriptions, mass photometry, tRNA cleavage assays, plaque assays and ChIP-seq experiments, as well as analyzed data alongside H.C.L. I.U. and H.C.L. analyzed ChIP-seq data with input from A.A.G.. S.K. and B.R. designed, performed, and analyzed microscopy experiments. A.H. developed custom code for image analysis. M.S. and S.K.A. constructed *E. coli* mutant strains. C.T., A.A.G., and H.C.L. measured the toxicity of OLD variants. Under the supervision of A.T.W., T.A.N. identified and characterized additional RecBCD-responsive systems, which were characterized by C.T. and A.A.G. in a temperature-sensitive *recBC* background. S.K.A. and G.S. provided RecBCD expertise and advised data interpretation. H.C.L. wrote the manuscript with contributions from S.K., I.U., and A.A.G. All authors edited and approved the final manuscript.

