## Supplementary Figures for "RecBCD Prevents Bacterial Autoimmunity by Clearing Self-DNA Ends Produced During Replication Termination"

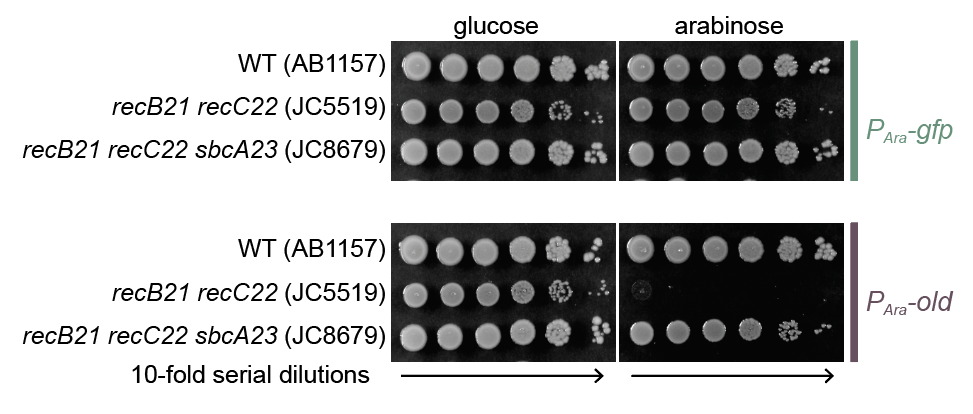

**Figure S1. *sbcA* suppresses OLD-mediated toxicity in a *recBC* mutant.**

Ten-fold serial dilutions of the indicated *E. coli* strains carrying plasmids encoding GFP (pNEB418) or OLD (pNEB37) under the control of *P_BAD_*. 5 µl dilutions were spotted on LB agar containing Cm15 and either glucose or arabinose, which represses or induces the production of the indicated proteins, respectively.

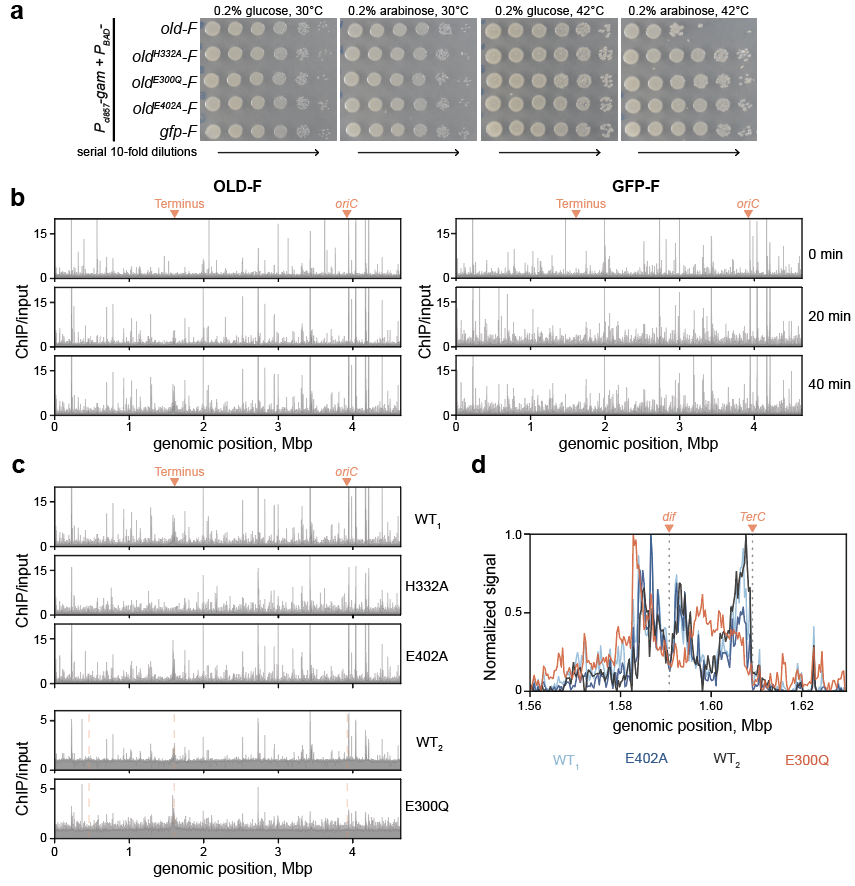

**Figure S2. ATP binding is necessary for DNA association and precedes tRNA cleavage**

(**a**) Ten-fold serial dilutions of wild-type *E. coli* carrying two plasmids: one encodes Gam under a heat-inducible promoter and the other encodes the indicated Flag-tagged proteins under the control of *P_BAD_*. The diluted cultures were spotted on LB agar containing Tet10, Cm15 and either glucose or arabinose to repress or induce FLAG-tagged protein production, respectively. Gam production was regulated by incubation at 30°C (repression) or 42°C (induction). (**b**) Input-normalized ChIP-seq profiles of OLD-F alongside the corresponding time-matched GFP-F controls. The data shown in Fig. 2c represent the same ChIP-seq datasets normlaized directly to these time-matched GFP-F controls. (**c**) Genome-wide view of ChIP-seq data presented in Fig. 2d. Data are shown relative to bacth-matched wild-type controls, with H332A and E402A compared to WT_1_, and E300Q was compared to WT_2_. (**d**) Min-max normalized enrichment profiles of the indicated samples. To highlight relative relative signal distribution, the ChIP/Input signals from Fig. 2d were background-substracted by subtracting 1.0 and scaled to the maximum signal within the shown genomic window. Data are representative of n=2.

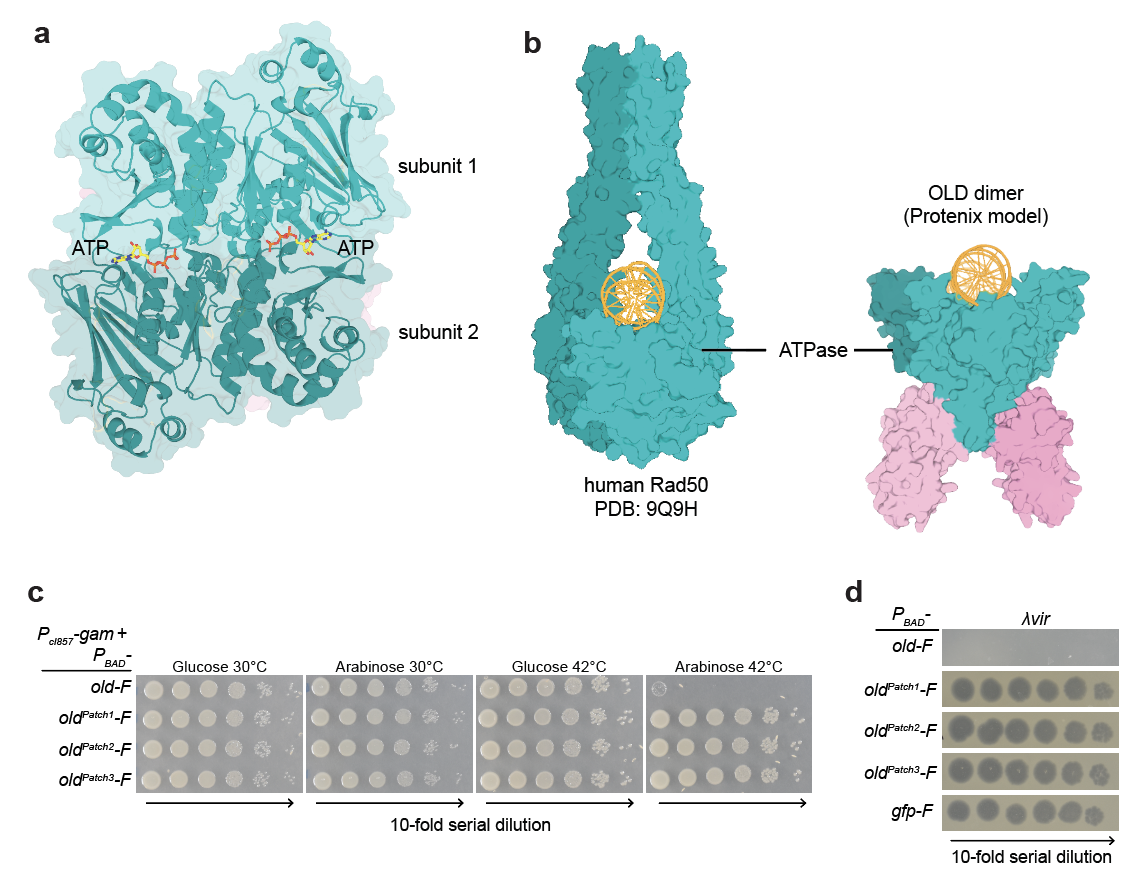

**Figure S3. ATP and positively charged surface faciliate phage defense and toxic function of OLD**

(**a**) Top view of the structural model of OLD dimer shown in Fig. 3a. Two ATP molecules (shown as sticks) engage the composite active sites formed at the interface between opposing ATPase domains of the OLD subunits. (**b**) Side-by-side comparison of human Rad50 dimer (left) and OLD dimer bound to DNA (right). The Rad50-DNA complex was derived from a cryo-EM structure of Rad50-Mre11 complex bound to DNA in which the Mre11 density was masked out, shown alongside a structural model of ATP-bound OLD dimers engaged with a DNA duplex. (**c**) Spot dilution assay to assess the toxicity of the indicated FLAG-tagged OLD variants. Experiments were performed as described in Extended Data Fig. 2a. Images are representative of n = 3. Quantification of this experiment was shown in Fig. 3e. (**d)** Plaque assay to assess defense against λvir. Ten-fold serial dilutions of λvir were spotted on lawns of wild-type *E. coli* producing the indicated FLAG-tagged OLD variants from a plasmid. See Methods for details.

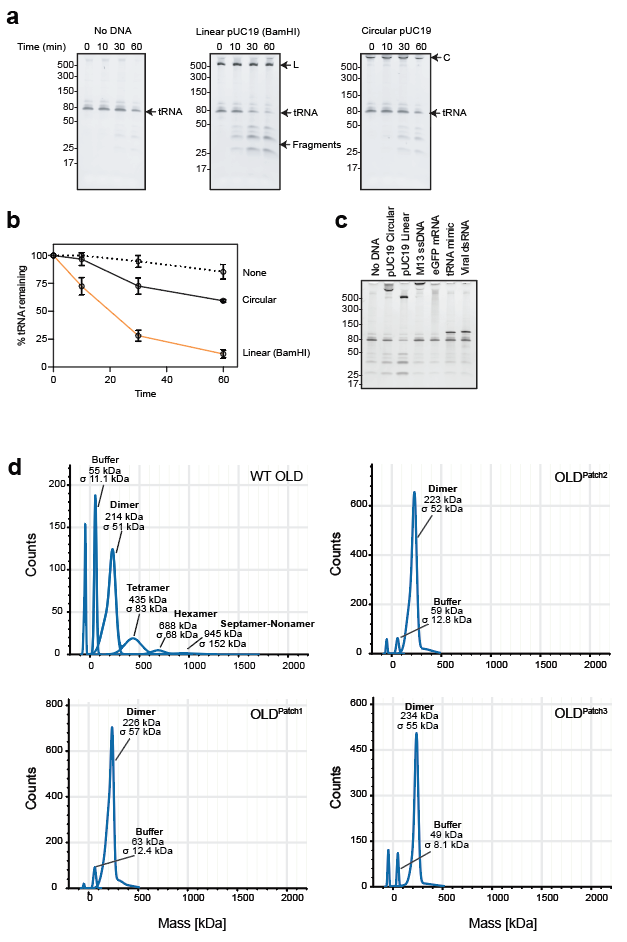

**Figure S4. OLD is activated by DNA ends**

(**a**) Gels showing cleavage of tRNA^ThrU^ by OLD over time in buffer containing 0.5 mM ATP and 250 mM K-Acetate. (**b**) Quantification of bands in (a). Data points are an average of triplicates and error bars depict standard error. (**c**) Cleavage of tRNA^ThrU^ by OLD in the presence of different nucleic acids. Reactions contain 50 ng of circular pUC19, linear pUC19 (BamHI linearized), M13 ssDNA, egfp mRNA, tRNA like structure of the Turnip yellow mosaic virus, or Adenovirus associated RNA1 as indicated, and were incubated for 30 minutes at 37°C. (**d**) Molecular weight distribution of 36 nM of the wild-type and the indicated OLD mutants in the presence of 1 mM ATP. Theoretical molecular weight of His-MBP-SUMO-OLD is 122 kDa.

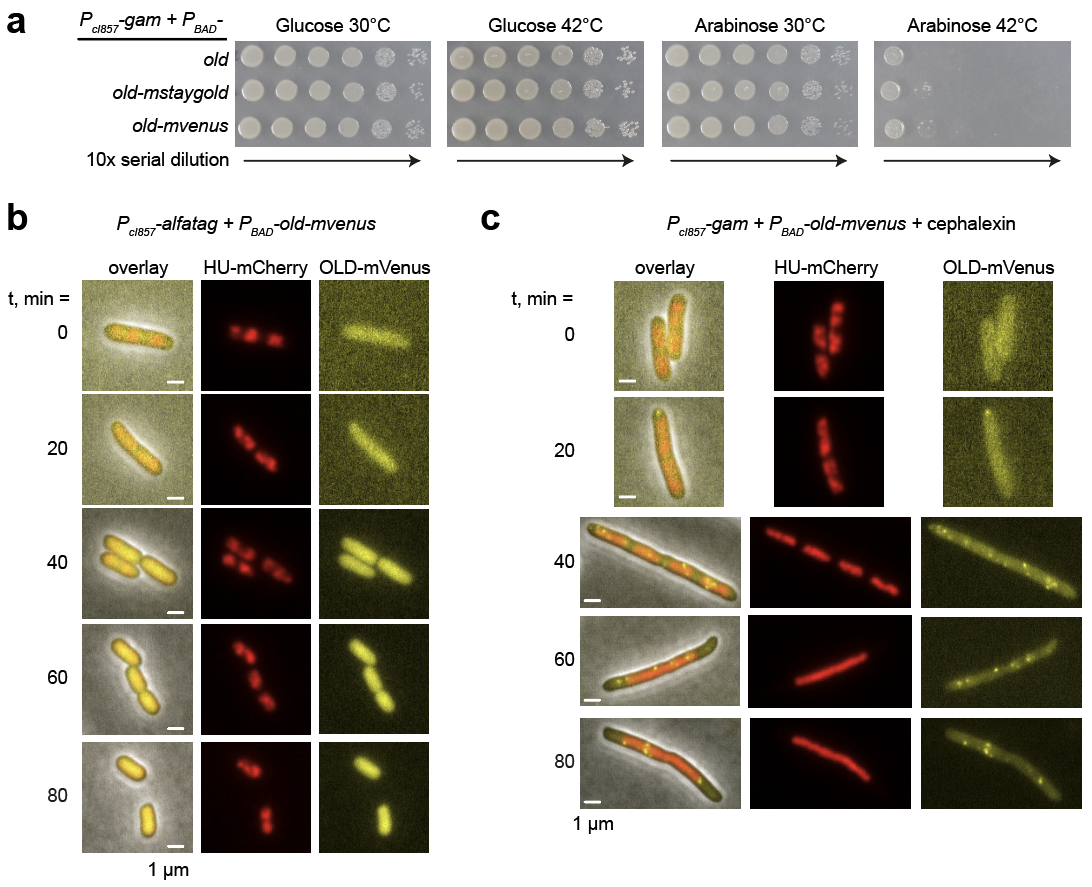

**Figure S5.** **Validation of OLD-mVenus focus formation.**

(**a**) Spot dilution assay to assess the functionality of fluorescent protein fusions. Experiments were performed as described in Extended Data Fig. 2a. (**b**) Representative time-course images of cells expressing HU-mCherry and OLD-mVenus following induction of the Alpha-tag. Columns show merged phase-contrast, HU-mCherry, and OLD-mVenus images (left), HU-mCherry fluorescence (middle), and OLD-mVenus fluorescence (right). OLD-mVenus remained predominantly diffuse, with no detectable focus formation over the 80-min time course. Scale bars, 1 μm. (**c**) Representative time-course images of cephalexin-treated cells expressing HU-mCherry and OLD-mVenus following induction of Gam. Cephalexin (50 ug/mL final) was added at the time of Gam induction. Images are displayed as in (b). Scale bars in (b–c), 1 μm.

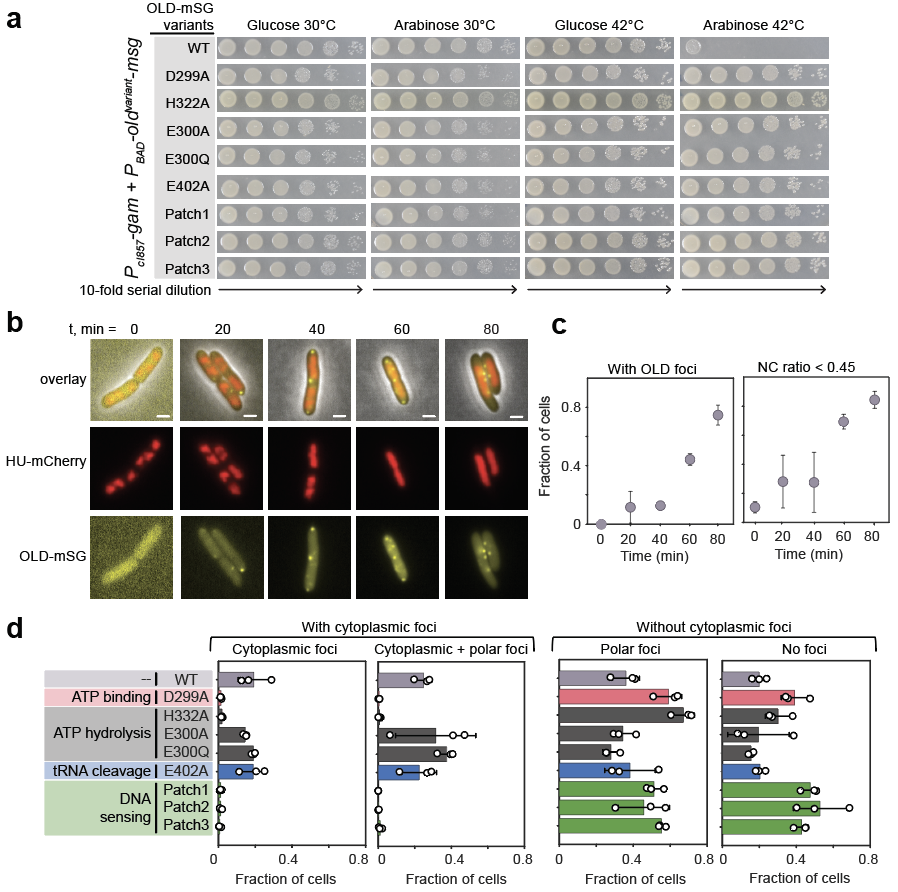

**Figure S6.** **Biochemical requirements for focus formation.**

**(a)** Spot dilution assay to assess the functionality of the indicated OLD variants fused to mStayGold (mSG). Experiments were performed as described in Extended Data Fig. 2a. Images are representative of biological n = 3. (**b**) Representative images of cells expressing HU-mCherry and OLD-mSG at the indicated times after Gam induction. Rows show merged phase-contrast, HU-mCherry, and OLD-mSG images (top), HU-mCherry fluorescence (middle), and OLD-mSG fluorescence (bottom). OLD-mSG exhibited progressive focus formation similar to that observed with OLD-mVenus. Scale bars, 1 μm. (**c**) Fraction of cells containing at least one OLD-mSG focus (left) and fraction exhibiting nucleoid compaction, defined as an NC ratio < 0.45 (right), at the indicated times after Gam induction. Results are shown as mean ± std across triplicates, including n = 505, 546, 1114, 1338, and 1019 cells at t = 0, 20, 40, 60, and 80 min, respectively. (**d**) Detailed classification of OLD–mSG localization for wild-type OLD and OLD mutants, corresponding to the data summarized in Fig. 4d. Cells were assigned to one of four mutually exclusive categories: cytoplasmic foci only, both cytoplasmic and polar foci, polar foci only, or no foci. The first two categories comprise cells with cytoplasmic foci, whereas the latter two comprise cells without cytoplasmic foci. Results are shown as mean ± std across three biological replicates. See Fig. 4d for data statistics.

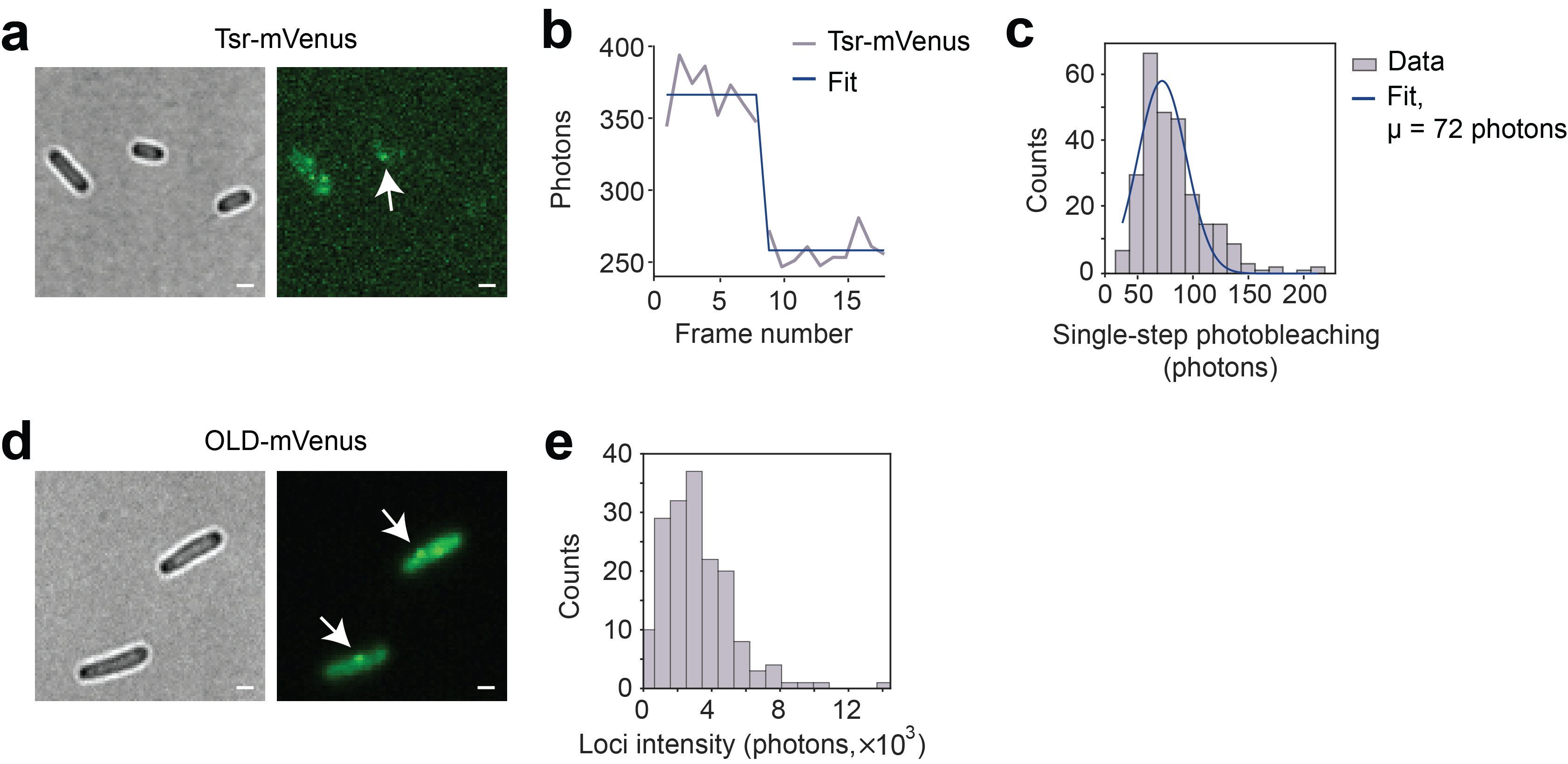

**Figure S7. Molecule count of OLD-mVenus per focus.**

(**a**) Representative phase-contrast (left) and mVenus fluorescence (right) images of cells expressing Tsr-mVenus. The arrow indicates a representative Tsr–mVenus fluorescent spot. Scale bars, 1 μm. (**b**) Fluorescence-intensity trajectory of a Tsr-mVenus spot under continuous illumination, showing a discrete photobleaching step. The measured trajectory (gray) was analyzed using a step-finding fit (blue) to determine the fluorescence-intensity change associated with individual photobleaching events. (**c**) Distribution of single-step photobleaching amplitudes measured from Tsr-mVenus spots. The solid line represents a Gaussian fit (μ = 72 photons and σ = 22 from n = 269 molecules). (**d**) Representative phase-contrast (left) and mVenus fluorescence (right) images of cells expressing OLD-mVenus at t = 80 min after Gam induction. Arrows indicate representative OLD-mVenus foci. Scale bars, 1 μm. (**e**) Distribution of background-corrected integrated fluorescence intensities of individual cytoplasmic OLD-mVenus foci (n = 179 foci), measured 60 and 80 min after Gam induction as shown in Fig. 3E.
